# Simulating human sleep spindle MEG and EEG from ion channel and circuit level dynamics

**DOI:** 10.1101/202606

**Authors:** B.Q. Rosen, G.P. Krishnan, P. Sanda, M. Komarov, T. Sejnowski, N. Rulkov, I. Ulbert, L. Eross, J. Madsen, O. Devinsky, W. Doyle, D. Fabo, S. Cash, M. Bazhenov, E. Halgren

## Abstract

**Background:** Although they form a unitary phenomenon, the relationship between extracranial M/EEG and transmembrane ion flows is understood only as a general principle rather than as a well-articulated and quantified causal chain.

**Method:** We present an integrated multiscale model, consisting of a neural simulation of thalamus and cortex during stage N2 sleep and a biophysical model projecting cortical current densities to M/EEG fields. Sleep spindles were generated through the interactions of local and distant network connections and intrinsic currents within thalamocortical circuits. 32,652 cortical neurons were mapped onto the cortical surface reconstructed from subjects’ MRI, interconnected based on geodesic distances, and scaled-up to current dipole densities based on laminar recordings in humans. MRIs were used to generate a quasi-static electromagnetic model enabling simulated cortical activity to be projected to the M/EEG sensors.

**Results:** The simulated M/EEG spindles were similar in amplitude and topography to empirical examples in the same subjects. Simulated spindles with more core-dominant activity were more MEG weighted.

**Comparison with Existing Methods:** Previous models lacked either spindle-generating thalamic neural dynamics or whole head biophysical modeling; the framework presented here is the first to simultaneously capture these disparate scales simultaneously.

**Conclusions:** This multiscale model provides a platform for the principled quantitative integration of existing information relevant to the generation of sleep spindles, and allows the implications of future findings to be explored. It provides a proof of principle for a methodological framework allowing large-scale integrative brain oscillations to be understood in terms of their underlying channels and synapses.

## 1. Introduction

Magnetoencephalography (MEG) and electroencephalography (EEG, together M/EEG), are complementary, non-invasive, instantaneous, and clinically essential, measures of human neural activity. M/EEG are measured as global brain activities, but are known to ultimately arise from channel currents, at a spatial scale ∼8 orders of magnitude smaller (Cohen, 2017). The causal chain which leads to M/EEG can be divided into two linked domains: (1) the biophysical propagation of electromagnetic fields after summating and cancelling under anatomical constraints; and (2) the neurobiology of large networks of active neurons whose ionic currents generate these fields. Here we present an initial effort to traverse the spatial scales by integrating simulations of large networks of neurons with biophysical models of electromagnetic propagation, informed by non-invasive imaging and invasive recordings in humans, as well as decades of work in animal models.

Ion movements through ligand- and voltage-gated transmembrane channels result in current flows which are influenced by the intrinsic channel properties of each neuron and the activity of the network. These currents flow through intracellular and extracellular spaces to form complete circuits, restricted by cellular membranes, and thus microscopic cellular anatomy (Einevoll et al., 2013). Currents cancel and summate locally with those of other neurons in the same cortical column, producing a net current which can be expressed as a multipole expansion (Nunez and Srinivasan, 2009). At a distance, the dipolar term predominates, and the local contribution is typically expressed as a current dipole moment. Before reaching the sensors, current dipole moments from different columns cancel and summate mesoscopically with other columns depending on their relative position and orientation in the highly folded cortical surface, and the covariance and phase synchrony of their magnitudes over time (Ahlfors et al., 2010a, 2010b; Irimia et al., 2012; Linden et al., 2011). Ultimately, the signal at each M/EEG sensor is the result of the complex cancellation and summation of microscopic synaptic and intrinsic currents from the many thousands or millions of neurons contributing to any single sensor’s leadfield.

Transmembrane currents are the result of spontaneous or evoked neural activity, which can be modeled computationally with various degrees of realism, balancing accuracy at the individual cell level against the quantity of neurons that comprise the simulated network. In the current study, we focus on a model for stage 2 of non-rapid eye movement sleep (N2) which is characterized by spontaneous sleep spindles. Sleep spindles manifest in normal M/EEG as spontaneous bursts of 10-16 Hz activity lasting 0.5-2 s and are thought to be important for memory consolidation (Andrillon et al., 2011; Bonjean et al., 2011; Contreras et al., 1996; Dehghani et al., 2011a; Diekelmann and Born, 2010; Sejnowski and Destexhe, 2000). A large number of studies in animal models have established the key elements in spindle generation: local circuit interactions between thalamocortical and thalamic reticular nucleus neurons, reinforcing intrinsic rhythmicity from successive activation of the hyperpolarization-activated cation current, I_h_ (Alain Destexhe et al., 1996; McCormick and Pape, 1990) and low-threshold Ca^2+^ current I_T_ (Huguenard and McCormick, 1992; Huguenard and Prince, 1992) Secondarily, the corticothalamic projections play a role in synchronizing and terminating the spindle (Bonjean et al., 2011; Timofeev et al., 2001).

Although the initial circuitry and cellular properties generating spindles are thus in the thalamus, the transmembrane currents that produce the M/EEG are cortical. The thalamocortical projection connecting these structures is comprised of a focal projection to layer 4 (termed the ‘core’), and a distributed projection to upper layers (termed the ‘matrix’) (Jones, 2002, 2001). We found previously that sleep spindles detected in MEG are more numerous and less synchronous than EEG spindles (Dehghani et al., 2011a, 2010), and suggested that this may reflect a relatively greater sensitivity of EEG to the matrix and MEG to the core projections (Piantoni et al., 2016). Consistent data has been obtained with laminar recordings showing primary involvement of middle versus upper layers in different spindles or parts of spindles (Hagler et al., 2018).

In this report we combine neural and biophysical models to generate M/EEG sleep spindles. The neural model is based on our previous computational modeling including the thalamic and cortical local and distant circuits involved in spindles, including matrix and core (Bazhenov et al., 2000; Bonjean et al., 2012; Krishnan et al., 2018b, 2016). All relevant thalamic ligand- and voltage-gated currents are included. The cortical elements are mapped to 20,484 locations on the ∼1 mm resolution cortical surface reconstructed from structural MRI. We have found in previous simulations that this resolution is necessary in order to accurately model the interactions between simultaneously active ECDs in producing M/EEG signals (Ahlfors et al., 2010b). In order to computationally model this large number of elements in cortex we use discrete-time models of neurons, which capture critical features of individual cell dynamics and synapses with difference equations (Rulkov et al., 2004; Rulkov and Bazhenov, 2008).

Empirical sleep M/EEG were collected to provide a basis for model evaluation and simulated neurons were embedded in donor cortical and cranial substrates produced from structural MRI collected from the same subjects. Thus, in our framework, it is the neural activity of individual persons that is modeled, as cortical geometry, since tissues intervening between brain and sensors, and connections among neurons are derived from the reconstructed cortical geometry of the subject. Microscopic cellular currents are scaled to mesoscopic current dipole moment densities using factors derived from human laminar electrode data. The extracranial electromagnetic fields generated by these mesoscopic sources are derived by quasi-static electromagnetic forward modeling, which accounts for orientation induced summation and cancelation by utilizing high-resolution cortical and cranial geometry. The basic validity of the model is tested by comparing the topography and amplitude of simulated macroscale extracranial M/EEG fields to those empirically recorded in the subject used to create the structural model.

The modeling approach employed here, an extension of our earlier work, allows for the currents of the coupled core and matrix networks to be isolated (Bonjean et al., 2012; Krishnan et al., 2018b) and then projected to the extracranial sensors (Gramfort et al., 2010). We find that simulated spindles have similar amplitudes and topographies to those recorded empirically, suggesting that the basic construction of the model is sound. We then apply the model to test the hypothesis that spindles recorded with MEG vs EEG tend to represent activity in the thalamocortical core vs matrix systems. Results consistent with that hypothesis are found. More generally, we demonstrate a proof-of-concept for relating microscale neuronal parameters to macroscale M/EEG observations.

## 2. Materials and Methods

The overall structure of the study is shown in Fig. 1. Two kinds of models were constructed: a Neural Model to compute sleep spindle activity during N2 sleep, based on the known anatomy and physiology of the thalamus and cortex; and a Biophysical Model to project the activity to the M/EEG sensors. Empirical Measurements were obtained and analyzed to produce Derived Measures, used to specify the models and validate the model: Structural MRI to define the location and orientation of cortical generating dipoles, Laminar recordings to scale the current dipole moment densities generating spindles, and M/EEG in the same subjects to permit validation of model predictions of amplitude and topography.

**Figure 1.**
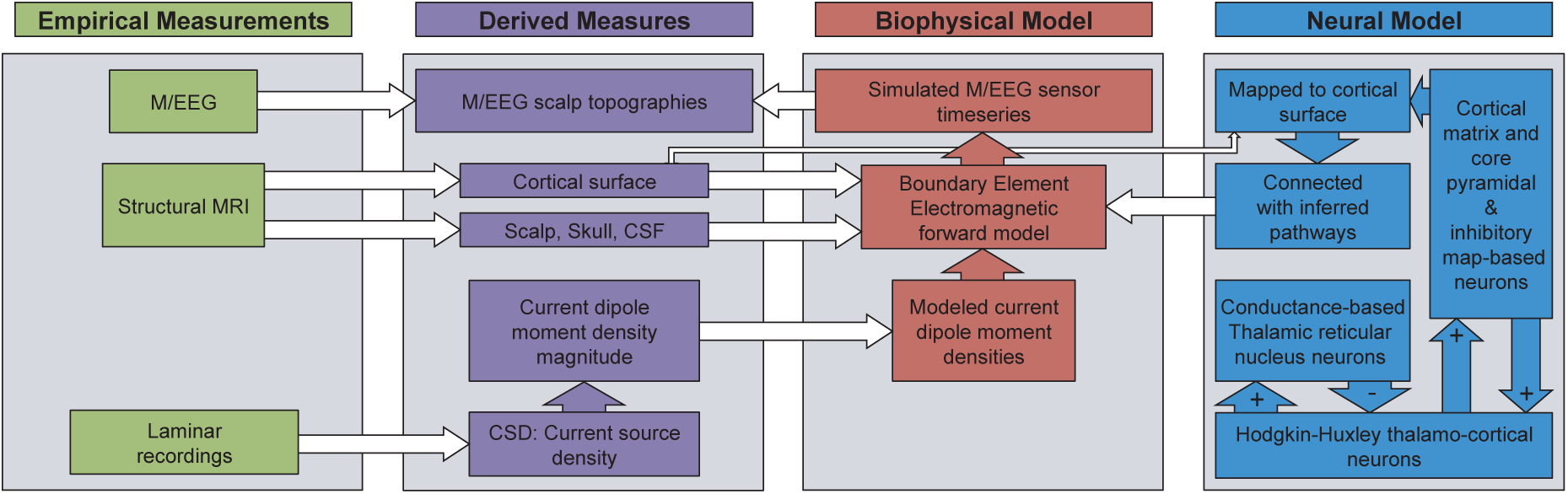
Overall structure of experiment. **Empirical Measurements** are processed to yield **Derived Measures** which provide validation targets (top), basic anatomical constraints informing the forward solution (middle), or basic physiological constraints for the fundamental unit of spindle generation (bottom). The **Neural Model** is comprised of thalamic cells modelled at high resolution (to capture the channel and local network synaptic processes underlying spindle generation) driving cortical cells (computationally-efficient map-based neurons), which are embedded in the cortical surface. The **Biophysical Model** takes the output of the Neural Model and projects it to the M/EEG sensors to be compared to actual empirical measurements.

### 2.1. Empirical Data

#### 2.1.1. Participants

MEG, EEG, and Structural MRI data were recorded for 6 healthy adults, (2 male, ages 20-35). Data for one additional subject was collected but was excluded from analysis due to poor EEG quality. Written informed consent approved by the institutional review boards of the University of California, San Diego or the Partners Healthcare Network, as appropriate, was obtained for all subjects. A whole-head MEG system with integrated EEG cap (Elekta Neuromag) was used to collect 204 planar gradiometers and 60 EEG channels. The position of the subjects’ head within the MEG helmet was monitored using head position indicator (HPI) coils (Uutela et al., 2001), updated every 15-20 minutes. Each subject’s headshape, HPI coil locations, and EEG electrode positions were digitized (Polhemus isotrak). Structural MR images were acquired in a separate session.

#### 2.1.2. M/EEG

M/EEG data were acquired during natural sleep at 1 kHz with a 300 Hz low-pass antialiasing filter. Epochs of stage II non-REM sleep were selected for analysis using standard criteria (Iber et al., 2007). Channels with poor data quality or gross artifacts were excluded by visual inspection. The electrocardiogram artifact was removed with independent component analysis (Delorme and Makeig, 2004).

#### 2.1.3. Structural MRI

High-resolution structural images were acquired with a 1.5 Signa HDx whole body scanner (General Electric). The acquisition protocol consisted of a 3-plane localizer, calibration scan, and a high-resolution T1-weighted MP-RAGE scans (TR = 10.728 s, TE = 4.872 ms, TI = 1000 ms, flip angle = 8 degrees, FOV = 256, 176 sagittal slices, 1 mm isotropic).

#### 2.1.4. Laminar recordings

As described in greater detail in (Hagler et al., 2018), after obtaining fully informed consent according to the Declaration of Helsinki guidelines as monitored by the local Institutional Review Boards, laminar microelectrodes arrays (Ulbert et al., 2001) were implanted into cortical tissue designated for resection in 5 patients (2 male; 15–42 years old) undergoing surgical treatment for drug resistant epilepsy. These arrays consisted of twenty-four 0.040 mm diameter 90%Pt/10%Ir contacts with 0.150 mm on-center spacing and were inserted along the surface normal of the cortex. Microelectrode localization within the cortical lamina was based on surgical procedure and electrode design and confirmed by histology in two patients. Bipolar referencing of reported laminar potentials yields a depth-resolved measure of potential gradients within and among cortical layers to be recorded simultaneously. After wideband (DC 10 kHz) preamplification (gain 10x, CMRR 90db, input impedance 1012 ohms), the laminar gradient recordings were antialiased at 0.5 kHz, gain 1000x, digitized at 2 kHz, 16 bit and stored continuously. Notch filters were applied to remove line noise and data from artifact-containing contacts were replaced by the weighted average of neighboring channels using an exponential decay function, λ = 0.1 channel spaces, (Hagler et al., 2018). Mild 1d spatial smoothing was applied with a Gaussian kernel (σ = 0.64) channel spaces, in order to ensure gradual and continuous variation across laminar channels, thereby suppressing false sources and sinks due to minor signal fluctuations.

### 2.2. Derived Measures

#### 2.2.1. Calculation of current dipole moment density scale

Periods of N2 sleep were isolated by the prevalence of generalized slow rhythms and spindles in simultaneously recoded cortical and scalp electrodes. Within these periods spindles were detected as described in Hagler et al. (2018). Briefly, after artifact rejection, spindles were identified as epochs with continuous sustained power in the 10-16 Hz spindle band (Andrillon et al., 2011). Spindle identification was made more selective by adding power in adjacent frequency bands as a rejection criteria (Mak-Mccully et al., 2017). Putative spindle epochs were detected independently for each laminar contact and epochs with durations less than 200 ms were rejected. Epochs containing a spindle in a least one laminar channel were identified and bounded by the earliest spindle onset and latest spindle offset across all channels, yielding a single unified set of detected spindle epochs for the entire array.

The laminar current source density (CSD), in µA/mm^3^, of sleep spindles was calculated by estimating the explicit quasi-electrostatic inverse of the laminar potential gradients (Pettersen et al., 2006). CSD’s were scaled by their distance from the center of the array to yield current dipole moments per unit volume, and then trapezoidally integrated over the length of the column to yield current dipole moment densities, in µAmm/mm^2^, or nAm/mm^2^. Microscopic transmembrane currents were scaled up to mesoscopic patch current dipole moment densities, a quantity corresponding to the dipole moment per unit area. This was accomplished by in two steps: scaling these currents by the cortical patch area represented by each column, then scaling these current densities to be consistent with empirical spindle current dipole moment densities recorded from human laminar microelectrode data

#### 2.2.2. M/EEG spindle topographies

Empirical and simulated M/EEG time series were band-passed to between 10 and 16 Hz with an 8^th^ order zero-phase IIR filter. The spindle-band complex analytic signal was extracted with the Hilbert transform and its envelope obtained by computing the elementwise modulus of the phasor time series. Spindles were automatically detected on empirical and simulated EEG standard criterion of sustained power in the 10-16 Hz spindle band (Andrillon et al., 2011). In short, the spindle band envelope was smoothed with a 300 ms Gaussian kernel (σ = 40 ms), and then normalized into units of standard deviation. Spindle occurrences were assigned to peaks of at least 2 s.d. and their temporal extent extended from these peaks until the smooth envelope fell below 1 s.d. For each detected spindle, the mean (unnormalized) envelope was computed and these data were interpolated over flattened sensor positions to produce topographic maps of spindle band envelope in a standardized head space (Oostenveld et al., 2011). Grand average maps (Fig. 6) were generated by averaging the mean spindle topographies from all subjects, or simulation runs.

#### 2.2.3. Core/Matrix index

The degree of core or matrix character was quantified for each simulated spindle. First, the 10-16 Hz envelopes were computed for the neural model derived current dipole moment density time series for core and matrix layers, using the procedure described for M/EEG analysis above. An index of core vs. matrix character was defined:

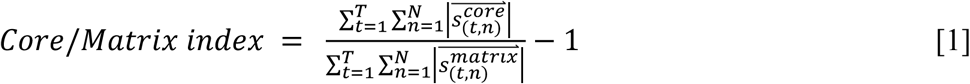

where *T* is the duration of the spindle in samples, *N* is the total number of cortical current dipole moment densities (20484), and 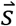 is the 10-16 Hz complex analytic signal. Positive values indicate a stronger core character and negative values indicate a stronger matrix character.

### 2.3. Biophysical Model

#### 2.3.1. Cortical reconstruction

White-gray matter surfaces were reconstructed from the MR volumes using FreeSurfer (Fischl, 2012). Surfaces were sampled at three resolutions as recursively subdivided icosahedra with 642, 10,242, and 163,842 vertices per hemisphere, wherein lower resolution meshes are nested subsets of higher resolution meshes. These are referred to as ico3, ico5, and ico7 surfaces, respectively, indicating the number of subdivisions. Within each hemisphere the geodesic distance between all pairs of vertices was computed using the Dijkstra approximation (Balasubramanian et al., 2009) with 13 Steiner nodes.

#### 2.3.2. Forward Model (Cortical current dipole moment densities to M/EEG)

The forward model, or gain matrix describing the gradiometer and EEG sensor activity produced by equivalent current dipoles at each ico5 vertex was then computed for each subject’s cortex. In addition to the white-gray matter surface, four extra-cortical boundary shells were reconstructed from the segmented (Fischl, 2012) images: gray-matter-cerebral-spinal fluid (CSF), CSF-inner skull, inner skull-outer skull, and outer skull-scalp. While the cranial boundaries consisted of triangular meshes with 5,124 vertices, critically, the cortical mesh was sampled at ∼1 mm resolution (327,684 vertices) in order to capture the summation and cancelation of opposed dipoles. The position of these surfaces relative to the EEG sensors and HPI coils (Uutela et al., 2001) was determined by registering the digitized headshape to the outer-scalp surface using non-linear optimization (matlab’s fmincon) with manual corrections. The position of these surfaces relative to the gradiometers was computed using the known relative positions between and the surfaces and the HPI coils, and the HPI coils and the gradiometers. The orientation of each dipole was set to the surface normal of the white-gray interface. The quasi-static electromagnetic forward solution was numerically computed using a four shell boundary element model, or BEM as implemented with the OpenMEEG software suite (Gramfort et al., 2010; Kybic et al., 2005). Consistent with reported experimental ranges, conductivities of 0.33, 1.79, 0.022, and 0.33 S/m, were used for the brain, CSF, skull, and scalp, respectively.

Rows of the resulting gain matrices were multiplied by the approximate Voronoi area (Meyer et al., 2003) of the cortical patch each represents to yield a vertex by sensor forward operator describing the contribution of each cortical patch’s current dipole moment density to each gradiometer and voltmeter. Current dipole moment densities resulting from core and matrix system pyramidal neurons were computed independently, summed together, and then multiplied by the forward operator to yield simulated EEG and MEG gradiometer time series.

Briefly, for the relatively low frequency of biologically relevant signals, electric and magnetic fields become uncoupled and the quasi-static approximations of the Maxwell equations can be used (Nunez and Srinivasan, 2009). Under this regime, the EEG forward model is a numeric solution for voltage, *V,* given *f*, the divergence of current density distribution, **J**^p^, in the Poisson equation:

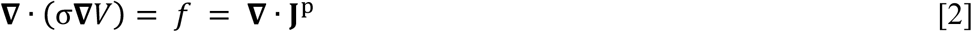

where σ is the tissue conductivity, in S/m. Because the cranial tissues are modeled as nested, closed, and piecewise homogenous domains, the integration reduces down to solving a symmetric linear system (Kybic et al., 2005). For MEG, solving for the magnetic field **B** requires both the current source distribution **J**^p^ and the computed electric field *V*, and is obtained by evaluating the Biot-Savart law at the boundaries:

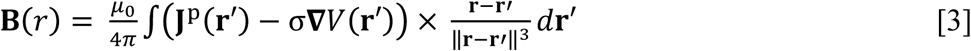

where **r** and **r**′ are displacements of the current source and magnetometer, respectively, and µ_0_ is the vacuum permeability constant. Planar gradiometer leadfields are derived by differentiating virtual magnetometer, or integration point, leadfields with respect to the length of the gradiometer. See (Gramfort et al., 2010; Kybic et al., 2005) for these methods in greater detail.

### 2.4 Neural Model

#### 2.4.1. Neurons

We used a computational model of a thalamocortical network (Fig 2A) with three layers in cortex, with each layer comprised of excitatory (PY) neurons and inhibitory (IN) neurons. The thalamus consisted of a network of core (specific) and matrix (non-specific) nuclei, each consisting of thalamic relay (TC) and reticular (RE) neurons. Conductance-based neural models were used to simulate thalamic neurons. A phenomenological model based on difference equation (map-based model) was used for cortical PY and IN cells. We have previously demonstrated that such map-based neurons are computationally efficient and able to reproduce the response characteristics of conductance-based neurons (Bazhenov et al., 2008; Rulkov et al., 2004; Rulkov and Bazhenov, 2008). Map-based models use a large discrete time step compared to the small integration time step used by pure conductance-based models while still capturing the dynamics of these models. Map-based models are capable of simulating large-scale network dynamics with emergent oscillatory activity (Rulkov et al., 2004) including slow oscillations during NREM sleep (Komarov et al., 2017).

**Figure 2.**
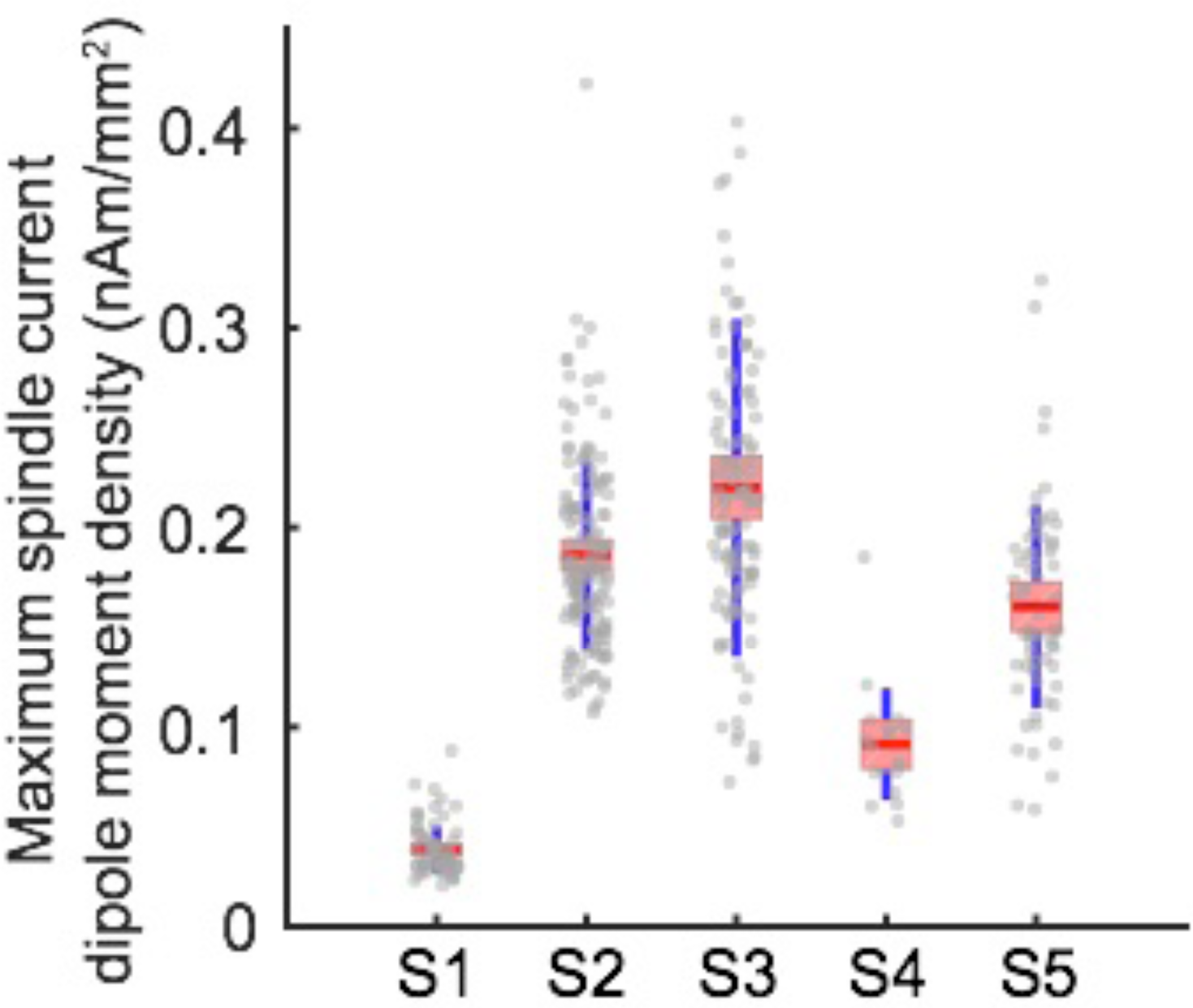
Dipole Magnitude of sleep spindles in humans. The current dipole moment density was estimated from CSD profiles as described in the text from laminar micro-electrode array recordings (24 contacts at 150µm centers, natural sleep). Gray dots represent the maximum current dipole moment density for each automatically detected 10-16 Hz spindle. Standard deviations and 95% confidence intervals are shown in blue and red, respectively.

<u>The following equation describes the update of the PY neurons in time:</u>

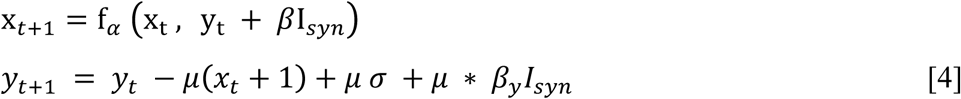

where variable x_*t*_ represents the membrane potential of a biological neuron at time t and y_t_ represent slow varying ion channel dynamics. The parameter μ (=0.0005) describes the change in the slow variable (μ less than 1 lead to slow update of y variable). The parameter *β* scale the input synaptic currents (I_*syn*_) for x variable, with *β* =0.133. The parameter σ (=0.02) defines the resting potential of the model neuron. The function f_*α*_ is given below:

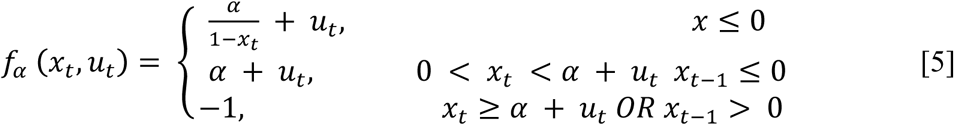

where *u*_*t*_ is taken as *y*_*t*_ + *β* I_*syn*_ from Eq 1, *α* (=3.65) is a control parameter which was set to obtain tonic spiking like activity for wide range of input currents.

<u>The inhibitory INs</u> were implemented using only the x variable to capture the fast spiking nature of inhibitory neurons and is described by the following equation:

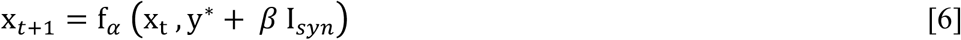

where, Y*=-2.90 with the same *f*_*α*_ function as Eq 2 with *α* =3.8 and *β* =0.05.

<u>The thalamic TC and RE cells were modeled as conductance-based neurons</u>, described by the following equation:

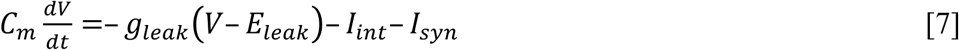

where the membrane capacitance, *C*_*m*_, is equal to 1 µF/cm^2^, non-specific (mixed Na^+^ and Cl^−^) leakage conductance, *g*_*leak*_, is equal to 0.0142 mS/cm^2^ for TC cells and 0.05 mS/cm^2^ for RE cells, and the reversal potential, *E*_*leak*_, is equal to −70 mV for TC cells and −77 mV for RE cells. *I*_I*nt*_ is the sum of active intrinsic currents, and *I*_*syn*_ is the sum of synaptic currents. The area of a RE cell and a TC cell was 1.43×10^−4^ cm^2^ and 2.9×10^−4^ cm^2^, respectively. RE and TC cells included fast sodium current, I_Na_, a fast potassium current, I_K_, a low-threshold Ca^2+^ current I_T_, and a potassium leak current, I_KL_ = g_KL_ (V - E_KL_), where E_KL_ = −95 mV. In addition, a hyperpolarization-activated cation current, I_h_, was also included in TC cells. For TC cells, the maximal conductances are g_K_ = 10 mS/cm^2^, g_Na_= 90 mS/cm^2^, g_T_= 2.2 mS/cm^2^, g_h_ = 0.017 mS/cm^2^, g_KL_ = 0.0142 mS/cm^2^. For RE cells, the maximal conductances are g_K_ = 10 mS/cm^2^, g_Na_ = 100 mS/cm^2^, g_T_ = 2.3 mS/cm^2^, g_leak_ = 0.005 mS/cm^2^. Fig. 3C shows a schematic illustration of the currents and synapses in our conductance-based neurons. The expressions of voltage- and Ca^2+^- dependent transition rates for all currents are given in (Bazhenov et al., 2002; Chen et al., 2012).

**Figure 3.**
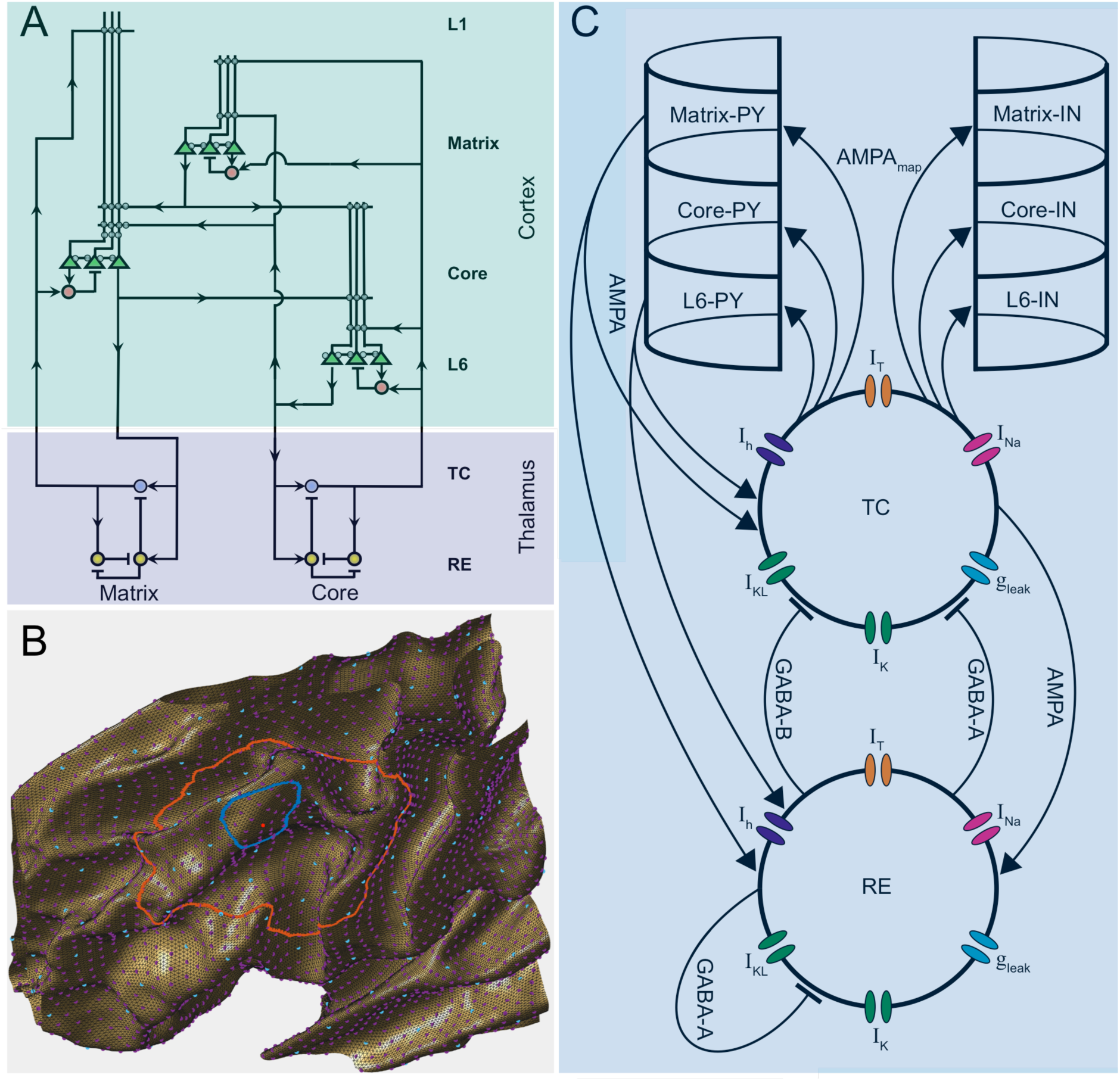
Network connectivity. (**A**) Schematic representation of thalamocortical and corticocortical connections. (**B**) Example of cortical geodesic-based connectivity in a patch of cortex. Pyramidal and inhibitory cortical neurons exist at purple and cyan locations, respectively. The blue contour shows the fanout (11.7 mm) of a core-projecting thalamic neuron at the virtual position marked in red. The orange contour shows the fanout (45.0 mm) for a matrix-projecting thalamic neuron at the same virtual location.(**C**) Schematic representation of currents and synapses included in detailed conductance-based thalamic neurons. Corticocortical connectivity is not shown. Please see text for details.

**Figure 4.**
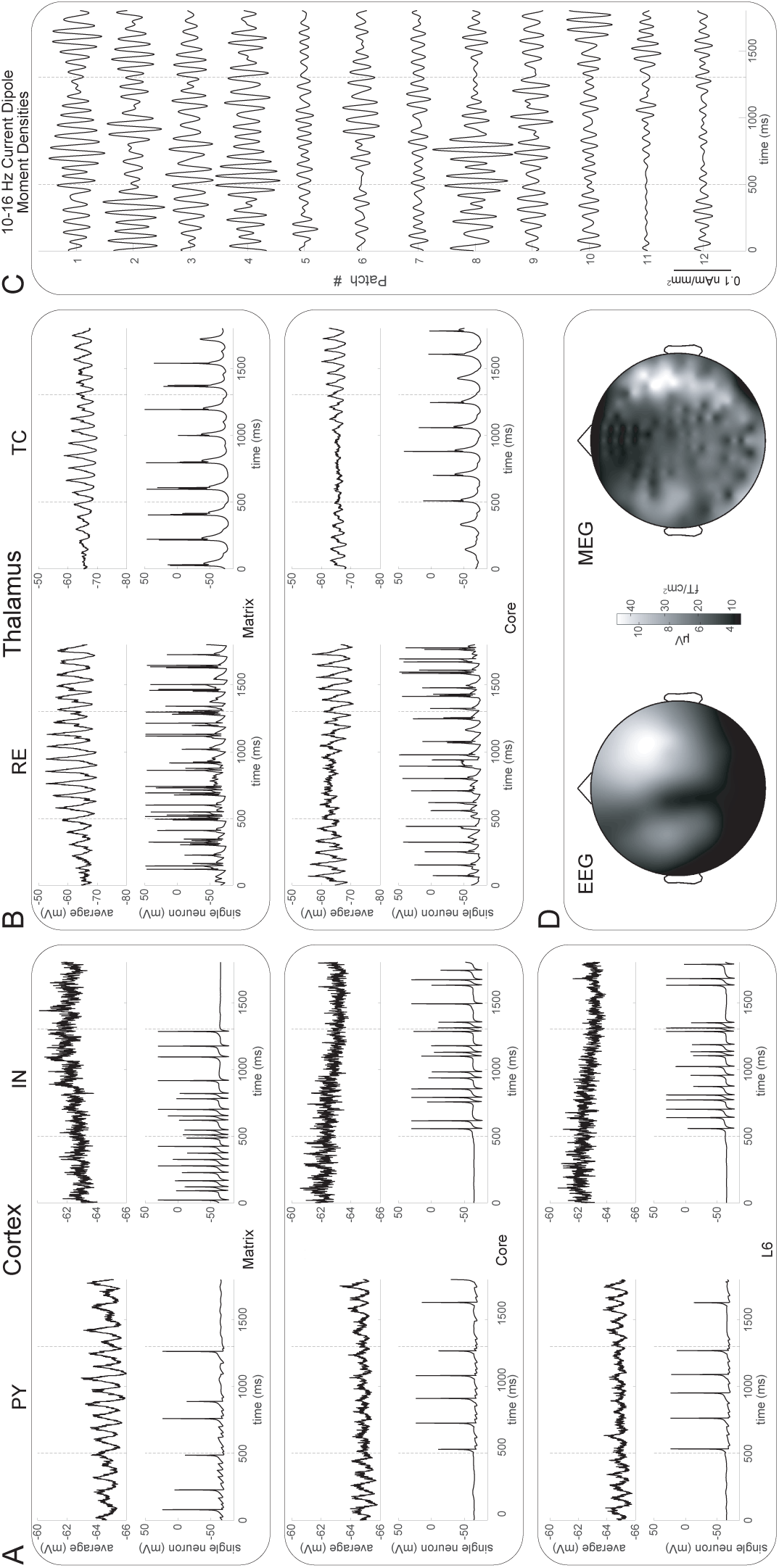
Example simulated spindle. Single-neuron and population mean membrane voltage traces in cortex (**A**) and thalamus (**B**) during a matrix-weighted spindle. Simultaneous activity is shown for the matrix and core system neurons of the reticular nucleus (RE) and thalamocortical (TC) subpopulations of the thalamus, as well as the pyramidal (PY) and inhibitory (IN) subpopulations in each of the three cortical layers. (**B**) Transcortical currents and scaled current dipole moment densities. Twelve spatially representative columns selected through icosahedral subsampling are shown. (**C**) Average 10-16 Hz M/EEG topographies for the duration of the spindle. In panels (**A**-**C**) the spindle duration is demarcated with vertical dashed lines. All panels display the same modeled spindle.

#### Synaptic currents

All the inputs to the map-based neurons were described by the

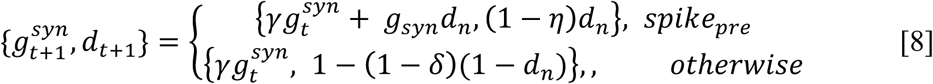

where 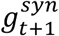 and *d* _*t*+1_ are the synaptic conductance and depression variable for time t+1, *g*_*syn*_is the synaptic coupling strength similar to the maximal conductance. The parameter *γ =* 0 99 is the decay variable to capture the first order kinetics of the synaptic transmission, *η =* 0.00005 is the rate of decay of the depression variable (d). The synaptic currents that are input to all conductance-based neurons were governed by equations given in (Timofeev et al., 2000) and reproduced here:

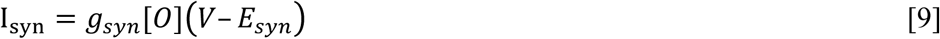

where *g*_*syn*_is the maximal conductance, [*0*] is the fraction of open channels, and E_syn_ is the reversal potential. In RE and PY cells, reversal potential was 0 mV for AMPA receptors, and −70 mV for GABA-A receptors. For TC cells, the reversal potential was - 80 mV for GABA-A receptors, and −95 mV for GABA-B receptors. GABAA, and AMPA synaptic currents were modeled by the first-order activation schemes. GABA-B receptors were modeled by a higher-order reaction scheme that considers the activation of K^+^ channels by G-proteins. The fraction of open channels [*0*] is calculated according to the kinetic equation:

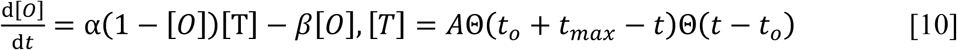

where Θ(x) is the Heaviside function, *t*_0_ is the time instant of receptor activation. The parameters for the neurotransmitter pulse were amplitude A = 0.5 and duration t_max_ = 0.3 ms. The rate constants, α and β, were α = 10 ms and β = 0.25 ms for GABA-A synapses and α = 0.94 ms and β = 0.18 ms for AMPA synapses. E was calculated according to the interactive scheme (Tsodyks and Markram, 1997).

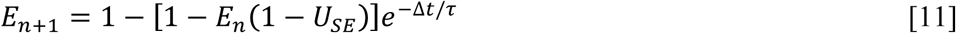

where Δt is the time interval between nth and (n+1)th spike, τ = 700 ms is the time constant of recovery of the synaptic resources and U_SE_ is the fractional decrease of synaptic resources after an action potential which was varied between 0.07 and 0.15. Spontaneous miniature EPSPs and IPSPs were included for the AMPA and GABA-A connections within cortical neurons. The arrival times of miniature EPSPs and IPSPs followed the Poisson process (Stevens, 1993), with time-dependent mean rate

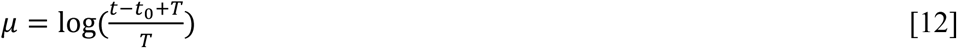

where *t* is current time and *t*_0_ was timing of the last presynaptic spike and *T* = 50 ms.

#### 2.4.3. Synaptic conductance

The maximal conductances for various connections were g_GABA-A_(RE-TC) = 0.045 μ S, g_GABA-B_(RE-TC) = 0.06 μ S, g_GABA-A_(RE-RE) = 0.175 μ S; core thalamus: g_AMPA_(TC-PY) = 0.03 μ S, g_AMPA_(TC-IN) = 0.015 μ S; matrix thalamus: g_AMPA_(TC-PY) = 0.045 μ S, g_AMPA_(TC-IN) = 0.02 μ S; connections within each layer (matrix, core and L6) pyramidal neurnons: g_AMPA_(PY-PY) = 2.5 nS, g_NMDA_(PY-PY) = 0.4 nS; connection from matrix to core: g_AMPA_(PY-PY) = 1.5 nS, g_NMDA_(PY-PY) = 0.1 nS; connection from matrix to L6: g_AMPA_(PY-PY) = 2.0 nS, gNMDA(PY-PY) = 0.2 nS; connection from core to matrix: g_AMPA_ (PY-PY) = 1.5 nS, g_NMDA_(PY-PY) = 0.1 nS; connection from core to L6: g_AMPA_(PY-PY) = 2.0 nS, g_NMDA_(PY-PY) = 0.2 nS; connection from L6 to matrix: g_AMPA_(PY-PY) = 2.0 nS, g_NMDA_(PY-PY) = 0.2 nS; connection from L6 to core: g_AMPA_(PY-PY) = 1.5 nS, gNMDA(PY-PY) = 0.1 nS; connection betwen PY and IN cells for all layers: g_AMPA_(PY-IN) = 0.05 μ S, g_NMDA_(PY-IN) = 0.4 nS, g_GABA-A_(IN-PY) = 0.05 μ S and connection from core and L6 cells to thalamic neurons: g_AMPA_(PY-TC) = 0.025 μ S, g_AMPA_(PY-RE) = 0.045 μ S.

#### 2.4.4. Network connectivity

A schematic of network connectivity is shown in Fig 2A. For each subject’s donor cortex, one pyramidal neuron was simulated for each vertex in the ico5 mesh (10,242 vertices per hemisphere) for each of layers matrix, core, and L6. The ico3 mesh was populated with inhibitory and thalamic neurons at each of 642 vertices per hemisphere. For all intra-hemispheric synapses, connectively was established by comparing synapse-specific fan-out radii to the geodesic distance between vertices on the ico7 cortex (163,842 vertices per hemisphere), see Fig. 3B. Inter-hemispheric synapses were threaded between homologously located cortical neurons in 85% of cases and the remaining connections were made between randomly located neurons. The probability of transmission in interhemispheric synapses was reduced to 25% and 50% in the core and matrix systems, respectively, in order to represent sparse collosal connectivity. Fig. 5B shows simulated current on an inflated cortex at a single time point for the core and matrix neurons.

**Figure 5.**
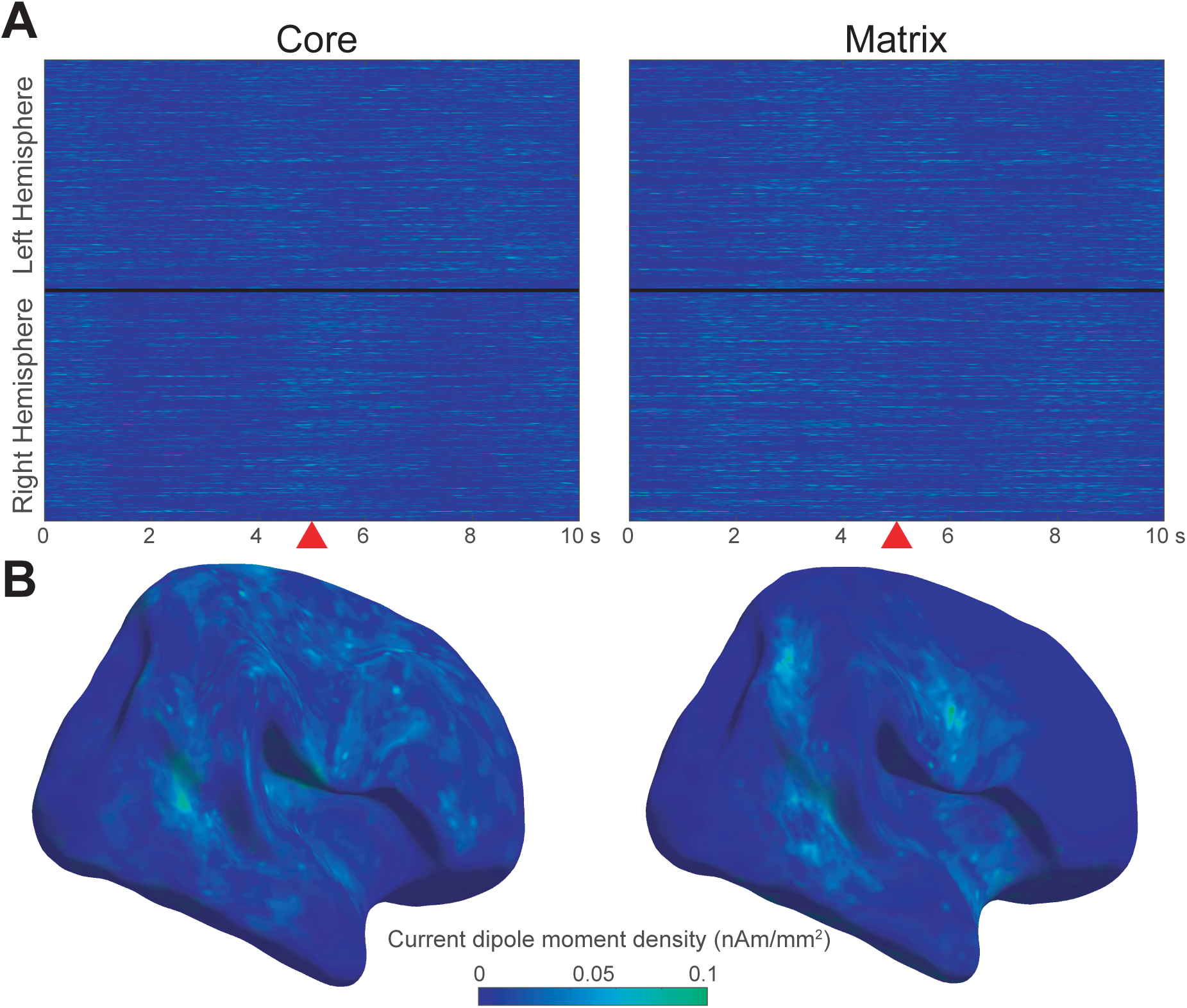
Simulated cortical sources. (**A**) 10 seconds of simulated spindling for one subject. (**B**) matrix and core pyramidal current dipole moment density distributed across the cortex at a single time point, marked in red. Data are displayed on an inflated right cortex.

## 3. Results

We designed a thalamocortical network model that combined the detailed laminar connectivity with the network connectivity of the whole brain based on MRI reconstructions. Using this approach we demonstrate the feasibility of connecting the cellular level activity with the macroscopic activity seen in M/EEG. We used a difference equation (map-based) model for cortical neurons, which has the computational efficiency necessary for simulating the cortex at sufficient resolution to accurately reproduce the cancellation and summation of cortical dipoles; we used conductance-based neuronal models for the thalamic network, which has the elements necessary to accurately reproduce the interaction of voltage-gated channels and recurrent synaptic connections central to spindle generation.

In a manner similar to our previous studies, (Bazhenov et al., 2000; Bonjean et al., 2011; Krishnan et al., 2016), the state of the network was set to be stage 2 sleep state by modifying the intrinsic and synaptic currents to mimic the level of low acetylcholine, nor-epinephrine and histamine. In this state, the network spontaneously generated electrical activity consisting of multiple randomly occurring spindle events involving thalamic and cortical neuronal populations. Spindle oscillations are driven by thalamic cell bursting as observed by experimental recordings (Steriade et al., 1993). Spindles spontaneously reappeared every 3–10s in agreement in prior intracellular data (Contreras et al., 1996) and computational models (Bazhenov et al., 2000; Bonjean et al., 2011; A. Destexhe et al., 1996; Destexhe et al., 1998). In this computational model, the initiation and termination of spindle sequences critically involved corticothalamic influences (Bonjean et al., 2011; Contreras et al., 1996; Timofeev et al., 2001). Furthermore, the synchronization of spindles across cortical and thalamic regions is determined by the strength and fanout of thalamocortical and corticothalamic connections (Bonjean et al., 2011; Krishnan et al., 2018b). The model consisted of two thalamocortical systems: core and matrix. The matrix system had broad thalamocortical and corticothalamic projections which is known to result in lower spindle density and increased spatial synchrony (Krishnan et al., 2018b). During spindles, cortical and thalamic neurons in both the core and matrix system had elevated and synchronized firing (Fig 3A) consistent with previous in-vivo experimental recordings (Steriade et al., 1993). This computational neural model was fed into a biophysical model: an electro- and magnetostatic forward model was applied to large-scale simulations of a thalamocortical network to simulate EEG and MEG signals.

As described in the Methods (2.2.1), current dipole moment densities were calculated using linear microelectrode array recordings spanning the cortical surface scale during sleep spindles in stage N2 sleep. The laminar current source density (CSD), in µA/mm^3^, of sleep spindles was calculated by estimating the explicit quasi-electrostatic inverse of the laminar potential gradients (Pettersen et al., 2006), followed by appropriate spatial scaling and integration over the cortical column to yield current dipole moment densities. As shown in Fig. 2, we found sleep spindle surface current densities have an average maximum spindle-band envelope on the order of 0.1 nAm/mm^2^ with considerable variation. Therefore, the simulated neural currents (in nA) were divided by the approximate Voronoi area (Meyer et al., 2003) of the cortical patch each represents, then scaled to approximately match in amplitude this surface current dipole moment density, yielding corresponding current dipole moment densities in nAm/mm^2^.

We found our model was able to simulate essential elements of empirical M/EEG. Grand average topographies of simulated and experimental data, shown in fig. 6, are qualitatively similar to experimental ones. The empirical MEG topography, in particular, is well-reproduced and shows the characteristic pattern (Dehghani et al., 2011b; Manshanden et al., 2002) of frontolateral gradiometer activation. The dynamic range across the scalp is higher in the simulated data, likely because the empirical data contains widespread non-spindle background activity, forming the 1/*f* curve of the power spectrum, and which was not included in the neural model.

**Figure 6.**
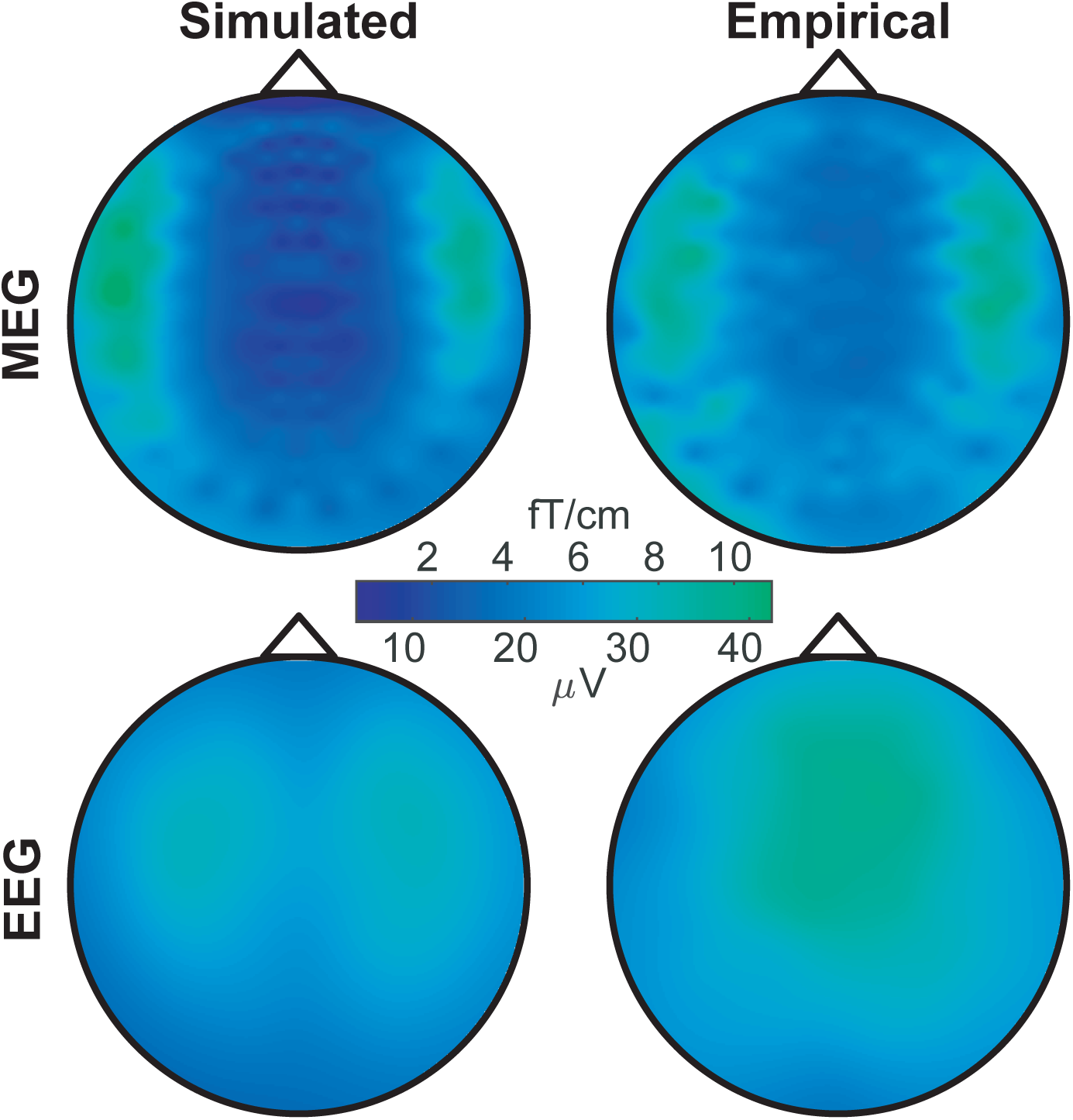
Grand average 10-16 Hz complex envelope topographies for empirical and simulated M/EEG during automatically detected EEG spindles. Empirical data show averages for six subjects, simulated data for six simulation runs using those subjects’ cortical surfaces, cranial tissue boundaries, and M/EEG sensor positions. The same MEG and EEG scales are used for both empirical and simulated topographies.

The simulated EEG topography, while matching the frontal position of the empirical data on the anterior-posterior axis, modestly differs in its lateral distribution. Whereas the empirical topography characteristically peaks along the midline, the two dorsal lobes of the simulated data only partially converge there. This may be due the model’s relatively crude implementation of inter-hemispheric connectivity, which consisted of low reliability synapses between mostly homologous cortical areas. An alternative possibility is that the ideal dipole model is incomplete for EEG (see below).

The simultaneously simulated MEG and EEG are also similar in magnitude of the empirical M/EEG (fig. 7A) with average spindle topography maxima (mean ± s.d. across all subjects or model runs) of 56.8 ± 12.6 fT/cm and 49.4 ± 13.4 fT/cm, p = 0.93, for simulated and empirical MEG, respectively, and 8.7 ± 0.8 µV and 10.7 ± 2.0 µV, p =0.96, for simulated and empirical EEG. However, despite their quantitative similarity, the simulated EEG and MEG spindles show opposite systematic tends, with simulated MEG spindles being slightly stronger and EEG spindles being weaker and less varied than empirical examples. This differential bias may be due insufficiently detailed or inaccurate cranial tissue conductivities, factors that have a much greater bearing on EEG than MEG, for whom these tissues are nearly transparent. Individual differences in skull conductivity (Akalin Acar and Makeig, 2013), in particular, may explain the increased inter-subject variability in empirical spindles. Another possibility is that the biophysical generation of these signals, commonly thought of as absolutely unified for a given source distribution, are in fact partially uncoupled, perhaps by the accumulation of static charges or effective monopoles, here unaccounted for, which would contribute to EEG but not MEG.

**Figure 7.**
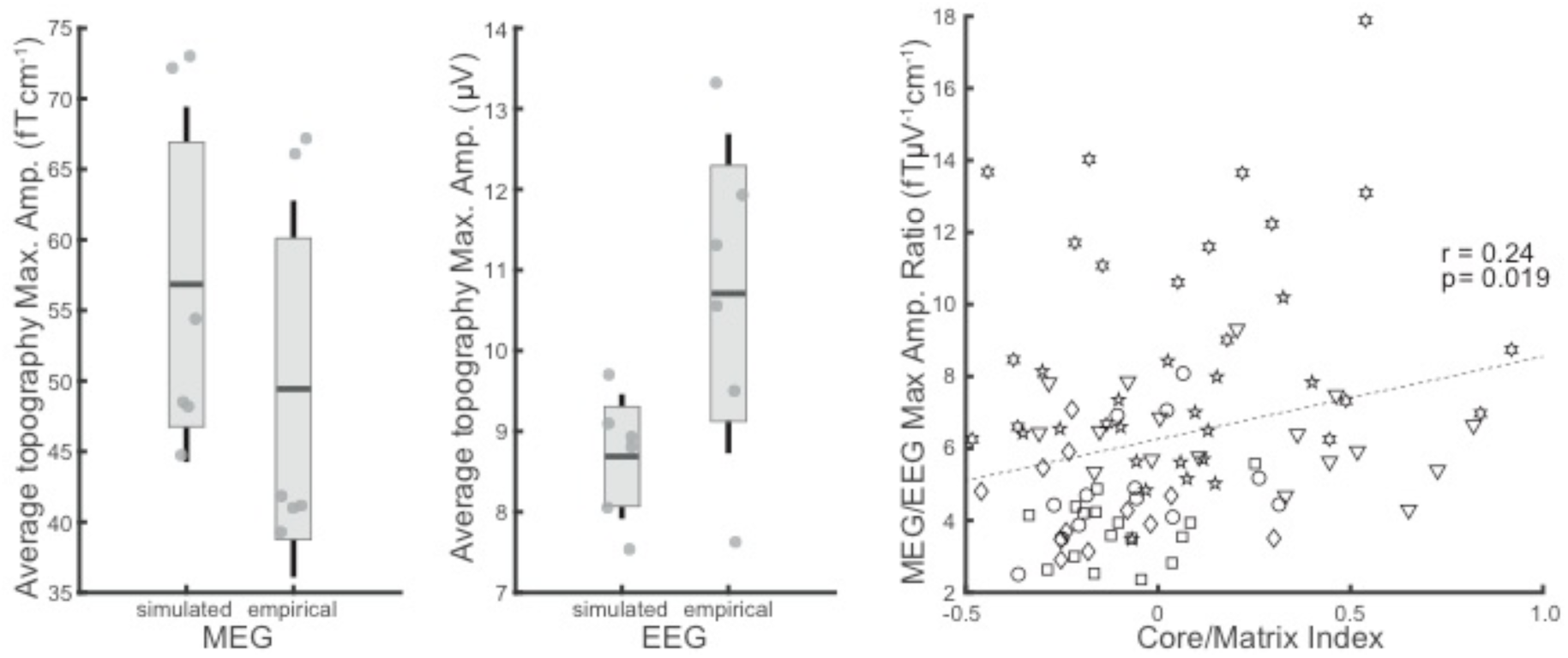
M/EEG sensor 10-16 Hz envelope maxima during EEG spindles. (**A**) For simulated and empirical M/EEG, 10-16 Hz envelopes are averaged across the duration of the spindles then averaged across spindles. The envelope magnitudes of strongest sensors are shown. Each marker represents the average value for one subject. (**B**) Simulated individual spindle sensor maxima vs. neural model derived core/matrix index. Positive index values indicate a more core-weighted spindle while negative values indicate a more matrix-weighted spindle. Each marker represents an individual simulated spindle with symbols signifying the simulation run, each using a different donor subjects’ cortical surfaces, cranial tissue boundaries, and M/EEG sensor positions.

For simulated spindles, the relative contributions of the core and matrix systems to current dipole moment densities was quantified and, as shown in fig. 7B, we found this index to correlate with the ratio of derived MEG vs. EEG maxima (Pearson’s r = 0.24, p = 0.019). These data are consistent with the hypothesis that MEG gradiometer recordings are more sensitive to core system neurons when compared to EEG recordings which are biased towards the matrix system. However, other factors, including individual differences, could also explain these results, and more focused studies are needed.

## 4. Discussion

In this study, we developed a computationally efficient large-scale hybrid thalamocortical model which generated sleep spindles in cortical patches. We estimated the magnitude of current dipole density from empirical measures and then used a biophysical model to project the output of the model simulation to the EEG and MEG sensors. The patches were embedded in the reconstructed cortical surface based on structural MRI data, and sufficiently dense to accurately model the summation and cancellation that occurs as M/EEG signals propagate from their cortical generators to the extracranial sensors. The amplitude and topography of the M/EEG derived from the model were similar to those found in empirical recordings in healthy subjects, when using their individual brain and head anatomy to define the projection from cortex to M/EEG, thereby suggesting that our approach is basically sound. We then applied our model to test the previous hypothesis (Bonjean et al., 2011; Dehghani et al., 2010) that EEG activity during spindles is relatively more sensitive to the matrix thalamocortical system, while MEG is relatively more sensitive to the core thalamocortical system.

Thus, using this approach, we demonstrate the viability of an integrated model for the generation of EEG and MEG, proceeding from ionic and synaptic activity, through local and distant networks, whose currents are then passed through a realistic biophysical model to generate M/EEG fields that correspond to empirical recordings, a foundational problem in neuroimaging. The model presented here simultaneously satisfies a considerable ensemble of neural and biophysical constraints including: dozens of neural properties, low-threshold Ca^2+^ current dynamics, the location and orientation of the cortical ribbon, empirically-backed biophysically probable maximum current dipole moment densities, and observed simultaneous M/EEG topographies and amplitudes. This multitude of constraints, embedded in each of the many scales the model traverses, precludes the pitfall of evoking over-tuned parameters in order to reproduce over-circumscribed behaviors. Given the ambitious scope of our model and the complexity of the system that it attempts to emulate, it is unsurprising that its limitations are numerous, including limited realism of the neural and biophysical models, limited application to M/EEG phenomena, and limited validation measures.

Our neural model simulated 65,304 cortical neurons and 5,136 thalamic neurons. While the model is thus much larger than previous efforts of this kind, it is still contains about 250,000 times fewer neurons than the actual human forebrain. Moreover, each simulated neuron is considerably simpler than a real neuron, especially in the number of synapses and number of dendritic compartments. In addition, subcortical areas other than the thalamus are not included in our model. While the direct contribution of these areas to extracranial M/EEG is minimal (Cohen et al., 2011), many, e.g. the nucleus basalis and hippocampus, may play critical roles in the timing, extent, amplitude and propagation of cortical activity during spindling. Notwithstanding these limitations, the model is sufficiently complex to generate sleep spindles using the same voltage-gated currents, within the same local thalamic and distant thalamo-cortico-thalamic synaptic circuits, as have been shown (Bonjean et al., 2012; Krishnan et al., 2018b) to generate sleep spindles in vivo. Furthermore, the model was sufficiently large to generate cortical patterns with a complexity that appears similar to that recorded in vivo (Frauscher et al., 2015; Mak-McCully et al., 2015; Piantoni et al., 2016) although this needs to be further investigated.

At its base, our study used a realistic computational model at the level of intrinsic and synaptic transmembrane currents to simulate MEG and EEG. It would be possible to increase the number of modeled neurons by using a population based neural mass model. Using such a model, Ritter et al. (2013) have modeled EEG signals from cortical activity projected to extracranial sensors. While such models could reproduce the spectrogram of EEG, they do not explicitly resolve activity at the level of individual neuron’s ionic or synaptic currents. This becomes critical when trying to leverage information from extensive intracellular and direct recordings of cortical and thalamic activity, to test new hypotheses. For example, abnormal spindling is common in schizophrenia (Wamsley et al., 2012) and variants of the gene CACNA1I, which encodes a T-type low-threshold Ca^2+^ channel and is expressed in the reticular nucleus of the thalamus (Manoach et al., 2016), are implicated in schizophrenia risk (Ripke et al., 2014). The framework we present can be used to simulate the effects of abnormal low-threshold Ca^2+^ channels on M/EEG. Costa and colleagues (Costa et al., 2016) included some of channel details in their neural mass model, but their biophysical modeling of EEG only considered an anatomically and physiologically implausible point source.

An important limitation of the neural model presented here is that it does not take into account functional specialization or cytoarchitectonic differentiation among cortical regions, including hemispheric lateralization. The model framework, however, can accommodate these distinctions. Projects using non-invasive structural, functional and diffusion MR imaging in large healthy populations now provide detailed cortical parcellations into distinct areas (Glasser et al., 2016). These can be combined with post-mortem transcriptomes to map receptor and channel variants, as well as laminar and cellular properties across the cortical mantle (Burt et al., 2017; Smith et al., 2013; van den Heuvel and Sporns, 2013). Future work could incorporate these cortical specializations into the neural model to determine if they underlie the variations in spindle amplitude and frequency across cortical areas which have been found with intracranial recordings (Frauscher et al., 2015; Mak-McCully et al., 2015; Piantoni et al., 2016).

Non-invasive imaging studies in humans and tracer studies in primates provide estimates of structural and functional connectivity between cortical areas (Glasser et al., 2016; Markov et al., 2014). The connections in our model did not incorporate these estimates but relied on the geodesic distance between cortical locations. While geodesics are a closer analog to anatomical and functional connectivity than conventional Euclidian distances, they remain a limiting simplification. A related limitation in our neural model is its lack of a conduction delay. Such delays are on the order of 10s of ms between lobes or between the thalamus and cortex (Klopp et al., 2000; Mak-Mccully et al., 2017), and thus, together with the strength and pattern of cortico-cortical connections could have a substantial effect on the propagation patterns (Muller et al., 2016), coherence, or phase relationships within and among spindles. These large-scale interactions are important for determining whether and how dipoles summate and propagate to the M/EEG sensors, and need to be addressed in future iterations of the model.

An additional limitation of our neural model is that the current dipole moment density produced by cortical spindles was based on empirical measurements, and the model only provided the timing, location and relative amplitude of the spindles. Current source density was calculated from sleep spindles recorded by 24 microcontacts spaced every 150 µm on center, traversing the cortical thickness from the pia to white matter (Hagler et al., 2018). The observed amplitude (∼0.1 nAm/mm^2^) is consistent with physiologically plausible maximum current dipole moment densities (Murakami and Okada, 2015). An important extension of the model would be to determine if this empirically determined value is consistent with that calculated in a detailed model of the columnar microstructure such as LFPy (Lindén et al., 2014) or the blue brain project (Markram, 2006). This calculation requires accurate reconstruction of dendritic domains, cell-densities and distributions, and synaptic terminations, in multi-compartment Hodgkin-Huxley models (Lindén et al., 2014). Such models are obviously too computationally expensive to incorporate directly into the many cortical patches in our model. However, an in-depth analysis of a single patch could help inform how well the limited number of cells we simulate in each patch represents the large number of actual cortical cells in that area.

The biophysical model used to propagate the cortical activity to the extracranial sensors also has limitations. We use a boundary element model which has been found to provide a good estimate of propagation at reasonable computational cost (Gramfort et al., 2010). Our model estimates four tissue-boundary ‘shells’ from each subject’s structural MRI: pia/CSF; CSF/skull; skull/scalp; and scalp/air. Although other models commonly omit the CSF layer, simulations indicate that its inclusion produces greater smearing of the EEG (Irimia et al., 2012; Lopes da Silva, 2013). Within the layers we use accepted tissue conductivities for the brain, CSF, and scalp (Hallez et al., 2007). However, the *in vivo* electroconductive properties of the skull remain controversial and may vary significantly across subjects (Akalin Acar and Makeig, 2013; Awada et al., 1998; Hallez et al., 2007). Furthermore, cranial nerve exits and other skull inhomogeneities may have significant effects (Akalin Acar and Makeig, 2013) which are unaccounted for in our model. Note that these issues will not affect MEG as, at the precision of biomedical analyses, the magnetic permeability of these tissues is equivalent to that of a vacuum (Hämäläinen et al., 1993).

The cortical surface used in the biophysical model is also reconstructed from each individual subject’s structural MRI (Dale et al., 1999). Extracranial M/EEG fields generated by a cortical dipole depend not only on its location and the magnitude of its moment, but are also highly dependent on the extent of spatial-temporal synchrony with other dipoles across the cortex and their relative orientations (Ahlfors et al., 2010a; Lutkenhoner, 2003). Dipoles are created by the post-synaptic currents of aligned pyramidal neurons oriented perpendicular to cortical surface (Lopes Da Silva, 2004; Nunez and Srinivasan, 2009). Thus, synchronous dipoles with opposed orientations, such as those on opposite side of the sulcal walls, will cancel each other out. In fact, the majority of the total cortical MEG signal is canceled before exiting the head on account of this phenomena (Ahlfors et al., 2010b; Lutkenhoner, 2003). Although the neural model contains only 20,484 cortical patches, their activity is mapped to 327,684 vertices for the biophysical model, which provides ∼1 mm resolution. Our previous simulations indicate that this resolution better captures the summation and cancelation of simultaneously active dipoles than lower resolutions (Ahlfors et al., 2010b). The high-resolution cortical mesh also reduces the numerical integration errors that can be present in BEM forward models with small inter-shell distances. However the efficacy of even finer resolutions at capturing cancelation accurately and the extent of errors in cortical ribbon orientation reconstruction are unknown, as are their effects on modeled M/EEG.

Despite these limitations of our neural and biophysical models, they produced reasonable amplitudes and topographies in both EEG and MEG. The amplitude of the M/EEG is a powerful constraint reflecting the interaction of many parameters and we are not aware of a previous study which reproduces both with realistic parameters and cortical source topographies. The empirical MEG topography, in particular, is well-reproduced and shows the characteristic pattern (Dehghani et al., 2011b; Manshanden et al., 2002) of frontolateral gradiometer activation. The leadfields of MEG gradiometers are smaller than those of EEG (Irimia et al., 2012), primarily because EEG is smeared by the skull and cranial tissues whereas these structures are mostly transparent to MEG (Hämäläinen and Ilmoniemi, 1994), especially when comparing bipolar MEG gradiometers to distantly referenced scalp EEG. Consequently, MEG is relatively more sensitive to focal sources whereas EEG to distributed (Irimia et al., 2012; Lopes da Silva, 2013). The core and matrix thalamo-cortical systems correspond to this pattern, with the core pathway terminating focally in layer 4 and the matrix more diffusely in superficial layers (Jones, 2002, 2001), leading to the hypothesis that core spindles would be relatively more prominent in MEG and matrix in EEG (Dehghani et al., 2010). We modeled the differential projections and terminations of the core and matrix systems and found support for this hypothesis.

In addition to the differing spatial extent of core and matrix spindles, their differing laminar distributions may also have an effect on their respective M/EEG signals. Laminar recordings in humans show that spindles can be dichotomized depending upon whether their LFP gradients are maximal in the middle versus superficial cortical layers, possibly corresponding to the core and matrix terminations of thalamocortical fibers (Hagler et al.,2018). The CSD calculated from the middle layer spindles produced a typical dipolar pattern with a current source and sink of approximately equal magnitude. However the superficial layer spindles consisted of a concentrated current sink and distributed or absent sources, yielding an effectively monopolar current distribution. Apparent monopoles are often found when analyzing CSD but it is controversial whether these represent an experimental or analytic artifact or a physiological phenomenon, such as accumulated charge (Bedard and Destexhe, 2013; Destexhe and Bedard, 2012; Gratiy et al., 2013; Riera et al., 2012). Most commonly, spindles are a mixture of middle and superficial layer waves (Hagler et al., 2018), and we did not distinguish between them when calculating the current dipole moment densities, which therefore mainly reflected middle layer spindles. Static ion concentrations produce an EEG but no MEG signal, and thus accounting for them in the matrix spindles in our model would have accentuated the difference between MEG and EEG in the predicted direction (i.e., producing a better correspondence between model and empirical results). Furthermore, the EEG signal produced by monopolar or unbalanced dipole decreases with distance less quickly than that produced by ideal dipoles, and this would tend to increase the similarity of the modeled to the empirical EEG spindle topography at midline sites.

Our model can be easily extended to other M/EEG phenomena such as K-complexes (Mak-McCully et al., 2014) and slow oscillations (Krishnan et al., 2018a; Wei et al., 2016) by marrying their neural computational models to the biophysical model described here. Using the same neural model to generate multiple M/EEG phenomena would provide a strong constraint on model parameters. Intracranial recordings may inform additional constraints, especially for the distribution, synchrony, and phase of cortical spindles (Halgren et al., 2018). Because the average spatial distribution of all spindles is much broader than that of individual spindles, the large-scale folding patterns of the cortex have a disproportionate impact on average spindle topographies. If simultaneous extra- and intra-cranial recordings are obtained, then the extracranial topography of individual spindles can be predicted by informing the model of the (limited) intracranial measures. This would permit predictions from individual spindles to be tested, rather than their grand average as was tested in the current paper.

A more comprehensive understanding of the relationship between neurobiology, local field potentials, and non-invasive M/EEG might plausibly improve the diagnostic power of the latter techniques. Here we have presented a framework that unifies detailed neural models with the measurement theory of M/EEG. Understanding the forward model that relates ion channel dynamics to M/EEG is the first step towards developing a principled inverse solution that maps M/EEG responses to clinically and physiologically relevant human *in vivo* molecular measures.

## Acknowledgements

This research was supported by NIH grants R01-EB009282, R01-MH099645-04, and T32-NS061847, as well as ONR grant N00014-16-1-2829. We thank Nima Dehghani, Donald Hagler, and Sergey Gratiy for their contributions to the empirical data, as well as our human subjects for their participation.

## Tables

**Table 1.** Method advantages and limitations

| Advantages | Limitations |
| --- | --- |
| Quantitatively relates neural and extracranial measures | Lacks detailed modeling of the columnar microstructure |
| Includes detailed dynamics of thalamic neurons including low-threshold $\text{Ca}^{2+}$ currents | Does not account for conduction delays in long distance connections |
| High resolution biophysical model captures effects of orientation and cancellation in cortical manifold | Example presented does not include areal diversity, but is possible within model framework |
| Can be use to probe intermodal differences and individual-specific anatomy and sensor configurations | Modeling of interhemispheric connectivity is rudimentary |
| Produces empirically observed M/EEG amplitudes from biologically plausible current dipole moment densities | White matter tractography not used to inform connectivity, planned for future iterations |

